# Mapping MAVE data for use in human genomics applications

**DOI:** 10.1101/2023.06.20.545702

**Authors:** Jeremy A. Arbesfeld, Estelle Y. Da, James S. Stevenson, Kori Kuzma, Anika Paul, Tierra Farris, Benjamin J. Capodanno, Sally B. Grindstaff, Kevin Riehle, Nuno Saraiva-Agostinho, Jordan F. Safer, Aleksandar Milosavljevic, Julia Foreman, Helen V. Firth, Sarah E. Hunt, Sumaiya Iqbal, Melissa S. Cline, Alan F. Rubin, Alex H. Wagner

## Abstract

The large-scale experimental measures of variant functional assays submitted to MaveDB have the potential to provide key information for resolving variants of uncertain significance, but the reporting of results relative to assayed sequence hinders their downstream utility. The Atlas of Variant Effects Alliance mapped multiplexed assays of variant effect data to human reference sequences, creating a robust set of machine-readable homology mappings. This method processed approximately 2.5 million protein and genomic variants in MaveDB, successfully mapping 98.61% of examined variants and disseminating data to resources such as the UCSC Genome Browser and Ensembl Variant Effect Predictor.

## Background

With the growing use of high-throughput sequencing technologies in the clinical setting, shortfalls in available genomic data to drive variant interpretation are increasingly observed. Specifically, clinical assessment of variants for pathogenicity are often inconclusive, with 48.79%(1) (1,372,553/2,813,113) of curated variants in ClinVar classified as “variants of uncertain significance” (VUS) at the time of writing due to the lack of clear evidence supporting or refuting pathogenicity(2). In silico prediction tools exist, but by themselves do not provide functional data that can help inform pathogenicity classification(3). In recent years, multiplexed assays of variant effect (MAVEs) have been introduced as a new line of functional evidence to support variant classification in a growing number of genes. MAVEs serve as useful tools for measuring the effects of variation on phenotype for thousands of variants in parallel(4). Commonly used MAVE designs include deep mutational scanning(5), in which the functional effects of protein variation in response to a selective pressure are described (6), and massively parallel reporter assays (MPRAs)(7), which validate the functions of different regulatory elements(8). As MAVEs can produce functional scores for many variants chosen systematically, they are able to generate functional evidence for variants of unknown significance (VUS) before they are detected in a clinical context, providing evidence that can ultimately assist in clinical variant interpretation (9)(10). MAVE data have already been incorporated into some ClinGen Expert Panel ACMG/AMP variant interpretation guidelines, e.g. for variants in *TP53* associated with Li-Fraumeni syndrome(11).

The increased use of MAVE experimental methods led to a need for central repositories for MAVE experiment metadata, assayed variants, and associated functional scores. In 2019, MaveDB(12) became the first such publicly-accessible resource. To improve data accessibility and discoverability, MaveDB organizes datasets in a hierarchical fashion using Experiment Sets, Experiments, and Score Sets. Experiment Sets are logical containers for linking multiple Experiment records, typically when multiple functional assays were performed on a single target as part of a study. Experiment records describe a single assay condition and its descriptive metadata, including the methods used and links to the raw sequence reads. Score Sets contain the score and optional count data for each variant measured in the assay and include details of the computational and statistical analysis performed as well as the target sequence. With nearly 300 submitted experimental datasets in MaveDB at the time of writing and more submitted every month, there is clear value in developing standards for the representation and exchange of these data, as well as standard methods for how these data may be calibrated and applied to support the clinical classification of genomic variants. The Atlas of Variant Effects Alliance (AVE; varianteffect.org) is a consortium working to realize these goals.

Under the auspices of the AVE Data Coordination and Dissemination (DCD) work stream, we have addressed the challenge of precisely mapping MAVE data to human reference sequences, and represent computational homology mappings using the Global Alliance for Genomics and Health (GA4GH) Variation Representation Specification (VRS) (13) and the Sequence Ontology (SO)(14) (**Figure 1**). For a given entry in MaveDB, a MAVE dataset describes sequence changes in MAVE-HGVS(15), a format similar to the Human Genome Variation Society (HGVS) variant nomenclature(16). MAVE-HGVS variants are then associated with their respective functional scores. These variants are represented with respect to a *target sequence* uploaded by the submitter; however, as this target sequence is not described with respect to human reference sequence sets (e.g. Ensembl/GENCODE(17), RefSeq(17,18), GRC genome assemblies), a challenge emerges concerning the standardized representation of variation. Specifically, while variants in HGVS are described in relation to accepted reference sequences (e.g. NP_003997.1:p.Trp24Cys), the vast majority of variants in MAVE score sets in MaveDB are described only in the context of the assayed sequence. For example, variant representations in MaveDB include “p.Ala40Ser” and “n.2G>A”, describing changes with respect to a provided *target sequence* stored in the MAVE experiment data (**Figure 2a**). Furthermore, the target sequences in MaveDB may not be reference-identical, may contain assay-specific functional elements that lack a comparable counterpart in the human genome, or align to different exonic regions of the human genome (**Figure 2b-d**). While necessary for the precise description of observed variants in an experimental setting, this design presents challenges to interoperability between MAVE datasets and variants described on human reference sequences, including variants routinely reported by clinical sequencing pipelines.

**Fig 1.**
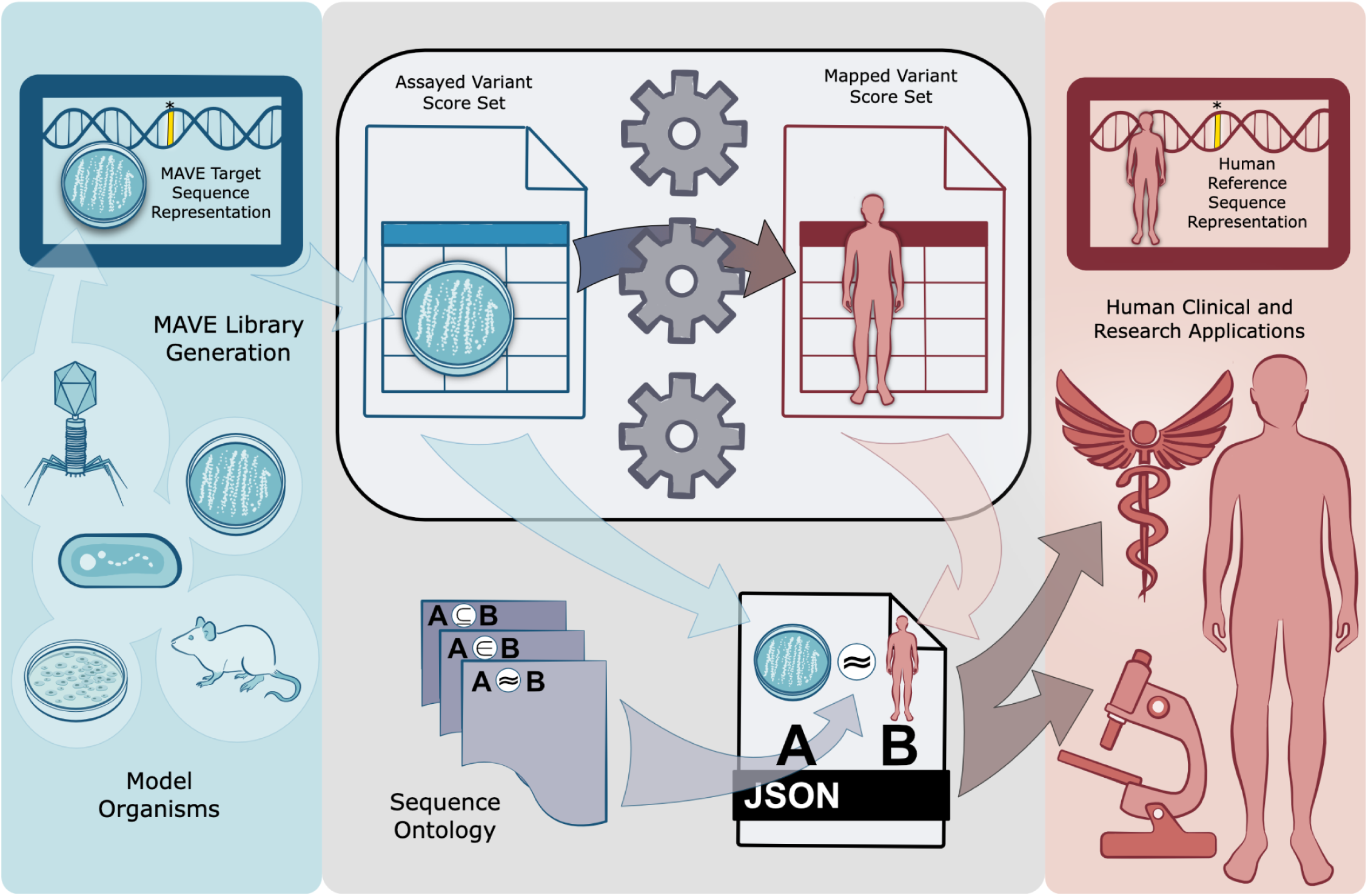
Mapping MAVE variants to the Human Genome. An overview of the MAVE variant mapping method. MAVE variants are described with respect to custom, user-submitted target sequences, but the absence of linkages to versioned human reference sequences limits the interoperability of MAVE data with human genomics applications (left). To overcome this limitation, we have developed a method to map MAVE variants to their corresponding human reference sequences (middle). Through the use of VRS, we are able to represent MAVE variants with respect to both assayed target sequences and versioned human reference sequences, creating robust homology maps (middle). The precise representation of MAVE variants using VRS ultimately facilitates the integration of MAVE data into downstream clinical and research applications (right).

**Fig 2.**
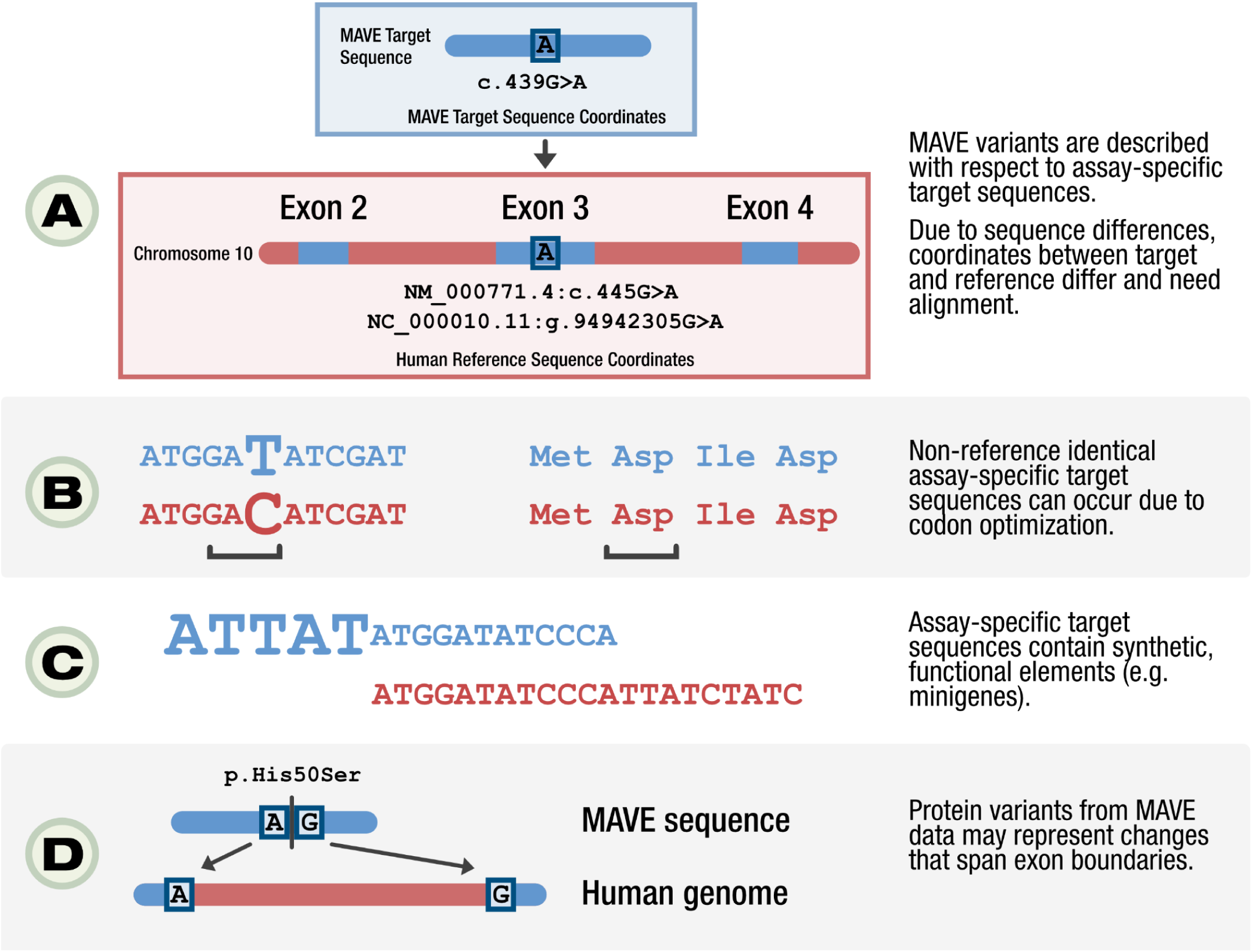
MAVE Assay-Specific Sequence Challenges. A depiction of several key features of MAVE sequences that necessitate a mapping strategy to human reference sequences. (**A**) MAVE variants are described using the MAVE-HGVS nomenclature system, which describes variants on a user-submitted target sequence. Therefore, as MAVE variants are described with respect to assay-specific target sequences, mapping to a human reference sequence is required to append an accession to each variant and add important contextual information. (**B**) MAVE target sequences are often non-reference identical due to features of the genetic system used in the assay. In the example above, there is a synonymous nucleotide substitution between the target and reference sequences, that optimizes translation of the sequence in the assay. (**C**) MAVE sequences can contain assay-specific functional elements that do not align to the human genome. (**D**) MAVE protein variants may represent changes that would span exon boundaries on the human genome, but occur on a contiguous region on the MAVE target sequence.

To resolve this limitation, we developed a method for consistently mapping MAVE variants to human reference sequences while preserving the original sequence context, improving provenance and interoperability between MaveDB and existing applications based on human reference sets. Using open-source tools and databases(13,19–21), we generated a MAVE dataset mapping for the FAIR(22) and computable exchange of variation data with other datasets and tools. We describe the integration of these mapped data into several common tools used for human genomics research and clinical variant curation, including the UCSC Genome Browser(23), the Ensembl Variant Effect Predictor (VEP)(24), the Broad Institute Genomics 2 Proteins Portal (G2P)(25), the ClinGen Data Platform(26), the DECIPHER resource(27), and Shariant(28). Our mapping approach closes an important gap for the application of MAVE data in genomic medicine and human health research.

## Results

### Composition of MaveDB Score Sets

Human score sets were selected for testing of the variant mapping functionality. At the time of writing, 209 of an available 299 score sets in the MaveDB had the *Homo sapiens* classification, together totaling 2,499,044 variants, providing a large and heterogeneous dataset upon which the mapping process could be developed and tested. Of the 209 selected score sets, 168 described protein coding elements while the remaining 41 covered regulatory and other noncoding elements, respectively. Among the 209 examined score sets, 176 contained DNA target sequences while the remaining 33 contained protein target sequences (**Figure 3a**). While the human score sets were designed to reveal insight into human biology, 60/159 unique experiments (37.74%) contained data from experiments that were conducted in genetic models not derived from human cells, including organisms such as yeast, mice, and bacteria (**Figure 3b** and **Supplemental Table 1**).

**Fig 3.**
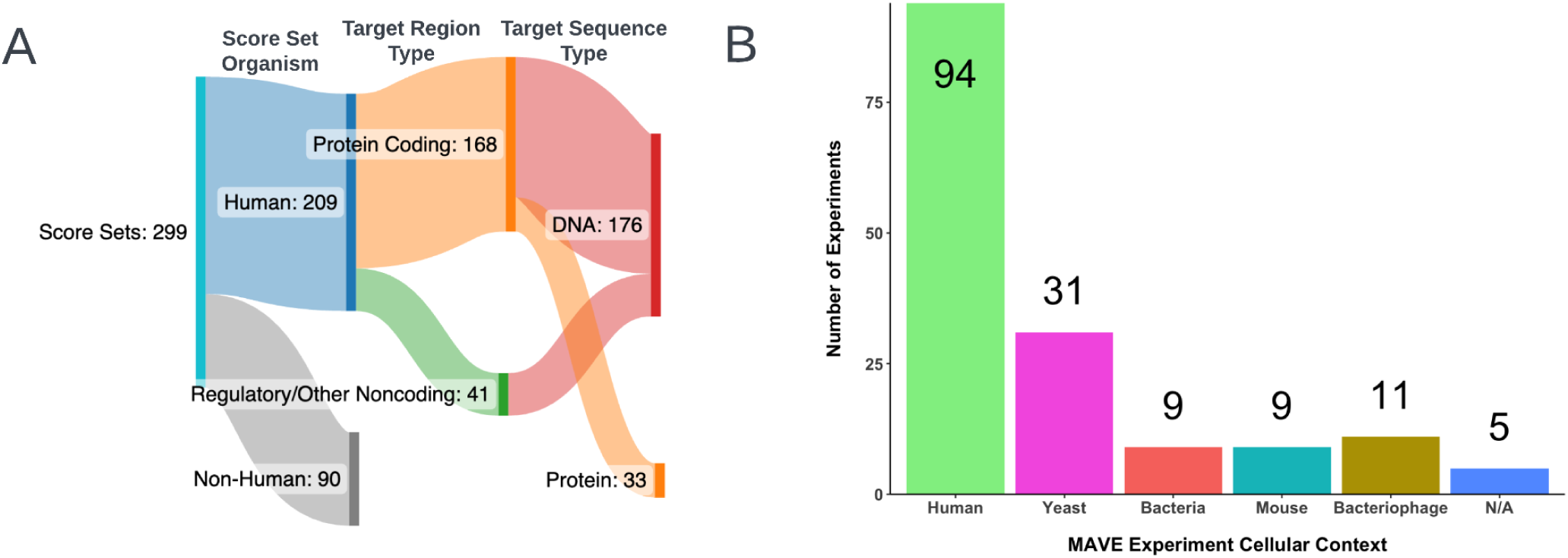
MaveDB Score Set Breakdown/Summary Statistics. A summary of the MAVE data from MaveDB that was used for validation of the mapping method. **(A)** All score set entries in MaveDB are assigned an organism attribute (e.g. *Homo sapiens*, *Saccharomyces cerevisiae*). Score sets whose listed target organism was *Homo sapiens* (n = 209) were selected for testing of the mapping algorithm, and additional breakdowns describing the selected human score sets are presented. Made with SankeyMATIC. **(B)** MAVE experiments in MaveDB (n = 159) can be conducted in non-human cellular contexts, including yeast, bacteria, mice, and bacteriophage(40), (41). Experiments that do not report a cellular context are coded as “N/A’’ (n = 5).

Most common among these experiment types were the 168 score sets describing protein coding variation, 95 of which reported associated UniProt(29) accessions. Variants from these score sets were described on either genomic (135 score sets) or protein (33 score sets) target sequences. The mean length of protein sequences (representing targeted regions of expressed human proteins) was approximately 573 amino acid residues, and the DNA target sequences had an average target sequence length of 1337 nucleotides. All 41 regulatory/other noncoding score sets reported DNA target sequences with an average length of 353 nucleotides. Gene symbols or aliases (e.g. *CALM1*, *CBS*) were provided for 126/168 protein coding score sets, and the remaining 42/168 reported specific domains and/or targets (e.g. hYAP65 WW domain, Src catalytic domain). All 41 regulatory/other noncoding category score sets described specific regulatory elements such as promoters (e.g *PKLR* promoter) or enhancers (e.g. *IRF6* enhancer). Across the 209 examined protein coding and regulatory score sets, 71 unique genes were included as score set targets.

### Mapping of MaveDB Score Set Variants to Human Reference Sequences

Variant mapping from human score sets was accomplished in three sequential steps (**Figure 4, Methods**). As homologous sequence annotations were not universally available across MAVE experiments, our first step was to use the BLAST-like Alignment Tool (BLAT)(21) to align MaveDB target sequences to the GRCh38 human genome assembly. Using this initial alignment data, we next computationally inferred compatible reference sequences (i.e. compatible RefSeq transcripts) associated with the score set, and the corresponding sequence offset for the MAVE target sequence. MAVE score set variants were then translated with respect to the associated RefSeq protein sequence or transcript-aligned genomic regions as appropriate (**Methods**) using the Biocommons SeqRepo(19) Python package/SQLite database and Universal Transcript Archive (UTA)(20) database and GenomicMedLab Common Operations on Lots of Sequences (Cool-Seq-Tool)(30) Python package. For score sets with regulatory/other noncoding elements, this second step was skipped, as the target could be directly mapped to a contiguous chromosomal region as aligned by BLAT.

**Fig 4.**
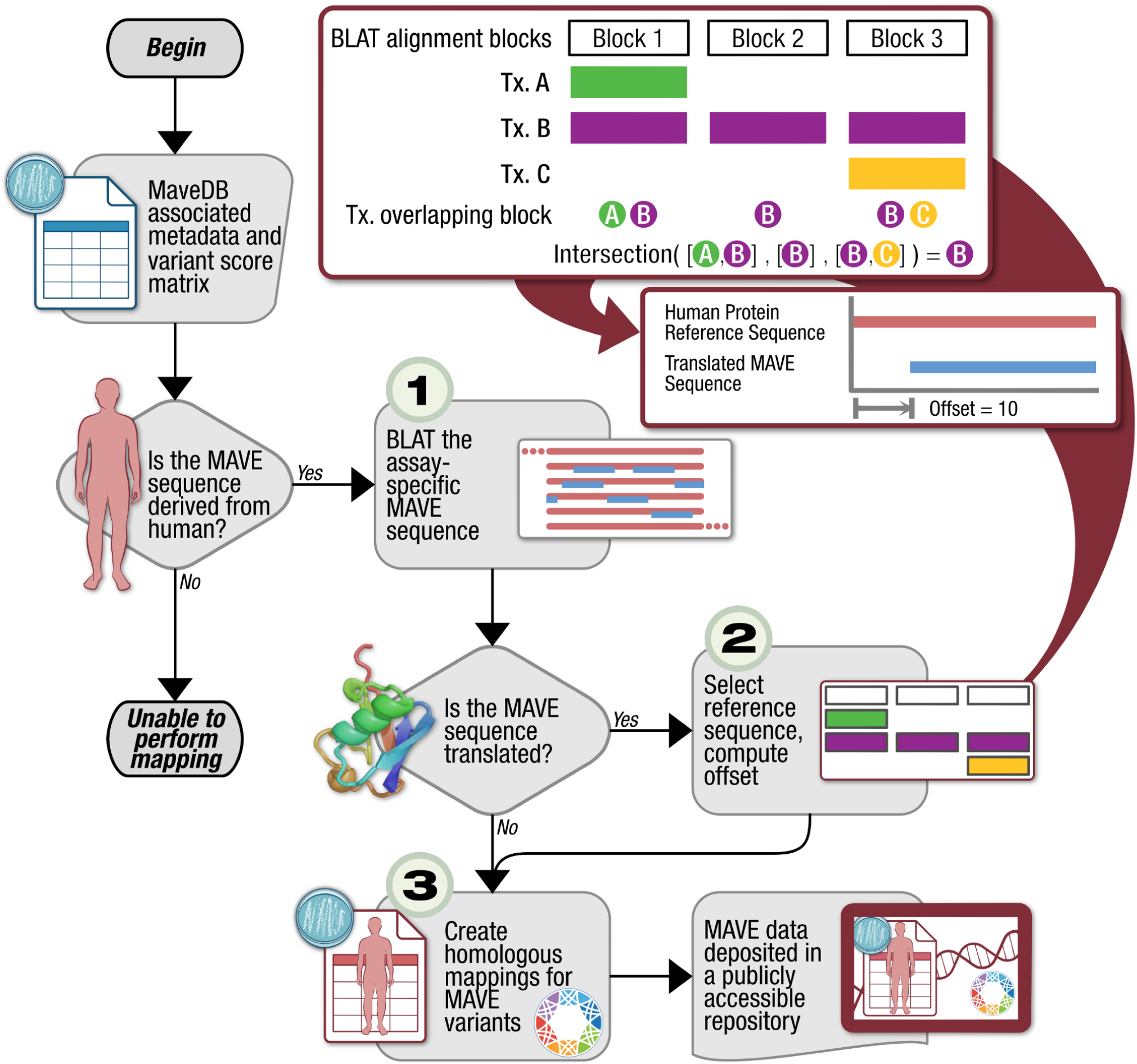
Variant Mapping Algorithm Workflow. A depiction of the MAVE variant mapping workflow. For a given entry in MaveDB whose listed target organism is *Homo sapiens*, the provided MAVE sequence is aligned to GRCh38 using BLAT(21), returning data including the chromosome number, gene symbol, and a set of genomic coordinates (**1**). If a score set describes a protein coding element, the outputted data can be supplied as a query to the Universal Transcript Archive (UTA) database, ultimately allowing for a RefSeq protein accession to be derived and for an offset to be computed (**2**). With a RefSeq sequence selected and offset calculated, the assayed variants in a MaveDB variant matrix are described with respect to their unique human reference sequence using the GA4GH Variation Representation Specification (VRS) (**3**). The resulting VRS objects are then annotated with descriptive metadata and integrated into specific score set JSON files. Lastly, the JSON files are gzipped and uploaded to a publicly-accessible s3 bucket to be available for downstream integration.

Lastly, the MAVE variants and mapped homologous variants were each translated into VRS alleles or haplotypes as necessary (**Figure 4**) and output along with the associated score data to comprise the resultant mapping set. Of the 209 human score sets that were potentially available for analysis, 207 were processed using the variant mapping algorithm. Of the two score sets that failed to map, the first (urn:mavedb:00000072-a-1) reported a UniProt accession that lacked any linkage to a corresponding RefSeq protein accession. The other score set (urn:mavedb:00000105-a-1) was unable to return a BLAT hit for the reference sequence. For the 207 human score sets in MaveDB, our method successfully mapped 98.61% (2,464,212/ 2,499,044) examined MAVE variants, where a successful mapping is defined as equivalence in the reference allele sequences of each pre-mapped and post-mapped variant pair. The remaining 1.39% (34,832/ 2,499,044) of discordant variants were caused by protein changes in the MAVE score set mapping across exon boundaries, non-homologous sequence content from MAVE target sequences preventing a reference match, and supplied variants occurring past the original target sequence length.

### Development of a Software Package for Mapping MaveDB Variants

To support continual MaveDB mapping efforts, we released our mapping pipeline as a Python software package. The three phases of the mapping workflow were constructed as separate modules, and additional methods were included to manage data acquisition from external sources. An included command-line interface enables end-to-end execution of the mapping workflow for a requested MaveDB score set, producing a JavaScript Object Notation (JSON) file that appends mapped scores to the score set metadata. The software was published to the Python Package Interface (PyPI) at: https://pypi.org/project/dcd-mapping/.

### Integrating Data from Mapped MaveDB Variants into Genomic Databases and Tools

#### MaveDB API

MaveDB includes a FastAPI-driven Application Programming Interface (API), specified at https://api.mavedb.org/docs. The API includes both pre- and post-mapped VRS objects for all variants mapped as part of this study, accessible as JSON using the MaveDB API `/mapped-variants` endpoint. These mappings are also available within the bulk downloads for mapped datasets. Future releases of MaveDB will further integrate native VRS support into the platform, by performing automatic mapping of submitted datasets using the protocols described here, and adding variant search capabilities using the contents of the mapped VRS objects. VRS objects will also power integration across different sources of genomic data as described here, and provide structured data for visualizations and other exploratory tools.

#### Genomics 2 Proteins Portal

Genomics 2 Proteins (https://g2p.broadinstitute.org/) Portal is an online discovery platform for linking genomic data to protein sequences and structures. The portal provides a user interface for exploring genetic variations, readouts from genetic perturbation assays, and protein features on the protein sequence and structure at the amino acid residue level to help interpret the molecular effect of variations. The portal integrates data from large genomic (gnomAD, ClinVar, and HGMD) and proteomic databases (including UniProt, PDB, and AlphaFold) as well as enabling users to perform customized mapping of genetic variations to proteins (**Figure 5a**).

**Fig 5.**
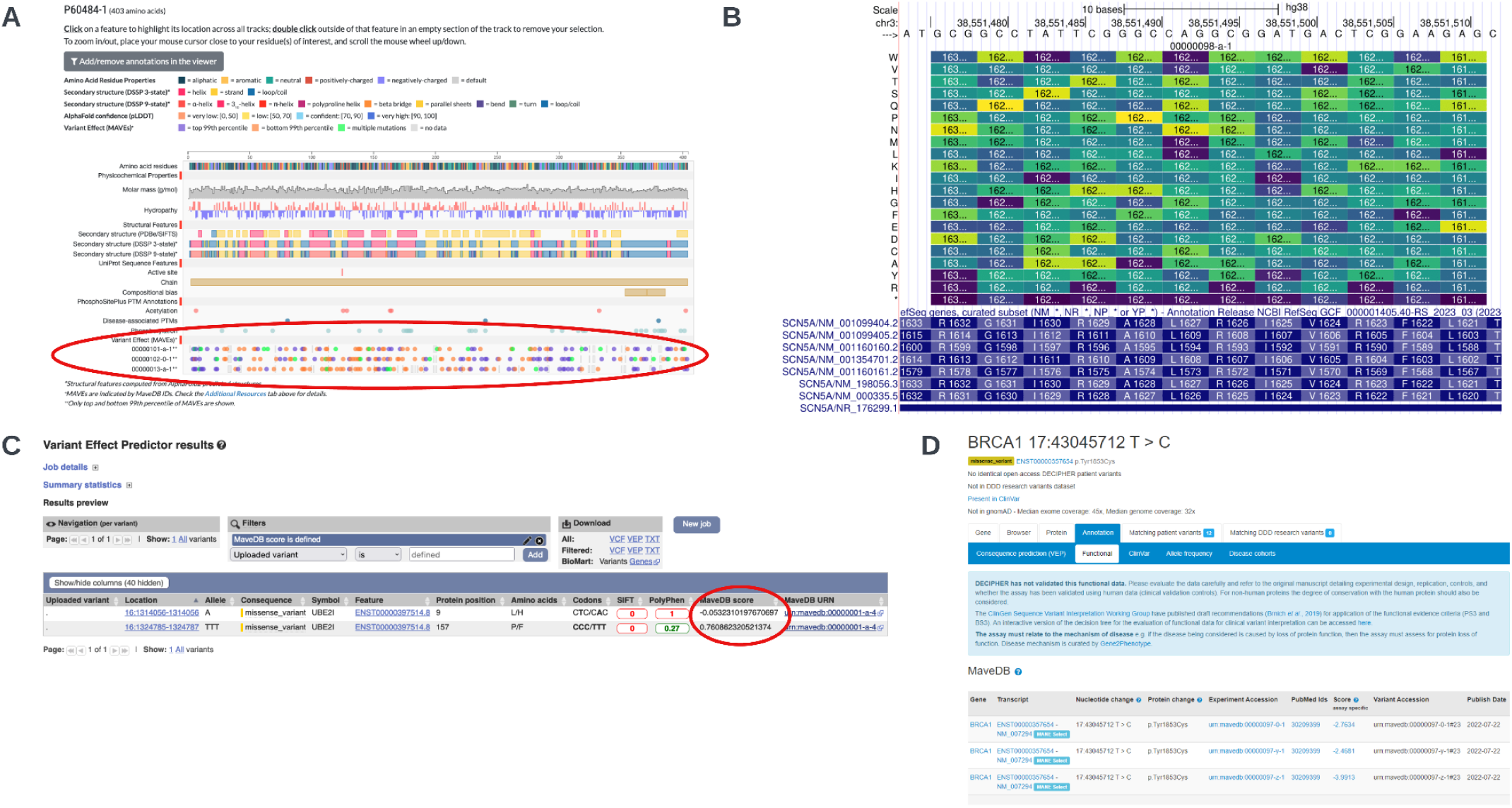
Downstream Integrations of MaveDB Data. The mapping of MAVE data to the human genome permits downstream data integrations in various human genomics applications. **(A)** MAVE scores are visible as heatmaps for available genes in the Genomics 2 Proteins Portal (located in the red circle). **(B)** MAVE data has been added as a track hub in the UCSC Genome Browser. MAVE protein variant positions are mapped to their corresponding genomic coordinates, and the score, chromosome band, genomic size, and strand are also reported for each variant. **(C)** MAVE scores and a link to the associated score set are reported, when available, for queried variants in the Ensembl Variant Effect Predictor (located in the red circle). **(D)** The nucleotide change, protein change, experiment accession, PubMed ID, assay-specific variant effect score, variant accession, and publish date are included for MAVE data displayed in DECIPHER, with links to the experimental details and score set in MaveDB. Example displayed: https://www.deciphergenomics.org/sequence-variant/17-43045712-T-C/annotation/functional.

168 score sets, describing MAVEs for protein coding variations in 40 unique human genes, have been integrated into the G2P portal. In addition to single amino acid residue substitutions (“point mutations”), MAVEs were available for pairwise residue mutations (“pairwise mutations”) for 7 out of 40 genes (**Supplementary Figure 1a**). While comparing to the length of the canonical protein isoform from Uniprot, MAVE data were mappable for >90% of the residues of 24 proteins (**Supplementary Figure 1b**).

All MAVEs for both point and pairwise mutations for each gene and score set are displayed in the G2P portal as heatmaps (**Supplementary Figure 1c-d**) and are downloadable as JSON files. Additionally, for each gene and score set pair, mutations with top and bottom 99th percentile of MAVEs were displayed in the context of protein sequence annotations of structural (secondary structure, residues’ solvent accessibility, etc.) and functional features (e.g., domain, active sites) (**Supplementary Figure 1e**). For example, MAVE readouts for *TP53* and score set urn:mavedb:00000068-b-1 range from −5.39 to 2.80. Mutations with scores >1.92 (top 99th percentile) and < −2.61 (bottom 99th percentile) were annotated in the “Protein sequence annotation” viewer of the portal. The filtering was performed for a clear visualization. These top and bottom 99th percentiles of MAVEs can also be mapped on their corresponding protein structure positions and are downloadable in tabular format from the portal. A list of genes with MAVE data in the G2P portal can be viewed under the “Protein Features” section on the statistics page at https://g2p.broadinstitute.org/stats. In future releases, average MAVE scores for each coding reference amino acid position will also be mapped to protein sequences and structures. The integration of MAVEs with protein sequence and structural features will facilitate interpreting MAVE data.

#### UCSC Genome Browser

The UCSC Genome Browser(23) is a widely-used and highly-customizable web tool supporting genome research, and includes annotations from many datasets of clinical and research relevance. A genome browser track hub has been created supported by these mappings, which displays the protein variant in genomic coordinates, and the associated scores. The track hub renders these scores as a heat map, in which each column represents the mapped genomic location of the variants scored, each row represents an alternate allele and cells are colored on a blue/yellow color spectrum in proportion to the score. This supports quick visual analysis to relate the functional impact of variants to their genomic context (**Figure 5b**). The MaveDB Genome Browser track hub can be accessed at https://genome.ucsc.edu/s/mcline/MaveDB, and under the UCSC Genome Browser Public Session gallery at https://genome.ucsc.edu/cgi-bin/hgPublicSessions. Each score set in MaveDB with mapped variants also includes a link to the associated mappings track in the UCSC Genome Browser for convenient access.

#### Ensembl VEP

Ensembl VEP (https://www.ensembl.org/vep) is an open-source tool for the annotation and prioritization of genomic variants. It aggregates currently available knowledge about variant loci and makes these available, with variant molecular consequence predictions, via three interfaces which have been designed to suit different use cases: 1) a highly configurable command line tool, 2) a REST API, and 3) a simple web interface. The genomic mapping of MaveDB data from this work has enabled the integration of MAVE score sets into Ensembl VEP. An extension to report these data has been developed for the command line tool, and we have updated the Ensembl VEP REST and web interfaces to annotate variants with information from open access MaveDB datasets (**Figure 5c**). This integration enables easy access to these data and convenient integration into large-scale variant annotation pipelines.

#### DECIPHER

DECIPHER (https://www.deciphergenomics.org/) is a global resource that shares phenotype linked variant data from rare disease patients to support research and diagnosis, and provides variant interpretation interfaces (31)(27). The mapping of MAVE data to reference sequence genomic coordinates has enabled DECIPHER to display this data in user interfaces, enhancing accessibility of this information to the clinical community. The MAVE data in DECIPHER are displayed on functional data tabs which are accessed from DECIPHER patient records in addition to variant pages and protein variant pages accessed through the site search tools (**Figure 5d**). Displaying these data in DECIPHER increases the discoverability of the MAVE data for clinicians, clinical scientists, clinical researchers, research scientists and curators who use DECIPHER, empowering its use in variant interpretation and assisting DECIPHER’s mission of mapping the clinically relevant elements of the human genome.

#### ClinGen Linked Data Hub

The ClinGen Linked Data Hub (LDH) is a RESTful API service built on Linked Open Data principles(32) that aggregates excerpts of pertinent variant data from a variety of external sources to contribute supporting evidence required for use variant curtation in the ClinGen Data Platform(26). The LDH works in concordance with the ClinGen Allele Registry(33) which is a canonical on-demand variant naming service. The mapped variants were submitted to the ClinGen Allele Registry and assigned Canonical Allele Identifiers (CAid) and Protein Allele Identifiers (PAid) for ingestion into the LDH.

Users can access the MaveDB data via the LDH API (note: UI also provides basic search functionality) by either using the LDH MaveDBMapping document’s entity ID (score set accession + “#” + variant number; e.g. urn:mavedb:00000001-a-1#1) or by searching for the associated variant CAid or PAid. Accessing the MaveDBMapping documents using the variant CAid or PAid allows users to easily access MaveDB data for the variant of interest from multiple MaveDB experiments or score sets simultaneously alongside pertinent data from other sources. The LDH API can also be used to return all MaveDBMapping documents from a particular score set, enabling bulk usage. Leveraging both ClinGen CAids / PAids and GA4GH VRS IDs allows for straightforward data aggregation of variants by identifier from groups that leverage one or multiple data standards and provides the users with the level of specificity required for their application. MaveDBMapping objects can be queried through LDH API (https://ldh.genome.network/ldh/MaveDBMapping/) and UI (https://ldh.clinicalgenome.org/ldh/ui/) endpoints (**Supplementary Figure 2**).

#### Shariant

Shariant(28) is a controlled-access platform to allow inter-laboratory automated sharing of clinically curated variants and structured evidence across Australian and New Zealand laboratories. The platform is configured to consume CAids from the ClinGen Allele Registry, which will be used to accomplish the initial data exchange between MaveDB and Shariant. This underscores the importance of integrating and supporting data standards, as the submission of VRS objects to the ClinGen Linked Data Hub is an essential step for generating the CAids that Shariant requires. MaveDB data linked to CAids and PAids will be made available to Shariant users as part of a planned platform update following user testing and feedback. Access to Shariant is restricted to Australian and New Zealand laboratories conducting clinical-grade testing.

## Discussion

In this study, we address the challenge of mapping multiplexed assays of variant effect (MAVE) data to human sequence assemblies for use in human research and clinical applications. We introduced and evaluated a method for accomplishing this using the GA4GH Variation Representation Specification (VRS)(13) and associated open-source bioinformatics tools. Our approach is informed by FAIR data principles and enables semantically-precise representation of these homology maps for data provenance.

However, our approach fails to map a small percentage of variants (1.39%; 34,832/ 2,499,044) due to a lack of policy about how such variants should be resolved. For example, when a DNA target sequence representing a processed RNA aligns to disparate regions of the genome, it is difficult to map insertions or deletions that span intron-exon boundaries, as the corresponding reference sequence genomic coordinates can cover thousands of nucleotides. Should these be treated as multiple insertion/deletion events, or as a single, very large event that also covers the intronic space? In other cases, segments of a target sequence may not align to known human reference sequences, limiting our ability to interpret variants that are reported at those unaligned positions of the target sequence. These are areas of ambiguity that would benefit from development of recommendations from the AVE expert community.

Our study also revealed diversity in the way variant knowledge is consumed and used by various downstream tools. GA4GH VRS provides a precise mechanism for addressing the complexity of representing variants on assay-specific sequences, and has useful characteristics for addressing variant overprecision and globally unique variant identification(13). However, its relatively recent emergence as a variant representation standard required additional mechanisms for enabling downstream connectivity to other resources, representing human-mapped variants as HGVS. To address this, we have used open-source translation tools to annotate all mapped variants using HGVS(34). We have also developed methods for mapping protein variants to the genomic reference space. Altogether, our approach has enabled integration of MAVE data into the Ensembl VEP, UCSC Genome Browser, Broad Institute Genomics 2 Proteins Portal, and ClinGen Linked Data Hub resources, with additional integrations forthcoming.

A remaining limitation for the applicability of these data to human clinical datasets is the development of expert guidelines for calibrating and scoring MAVE assays, work that is ongoing in the context of AVE. The responsible use of this data for clinical purposes relies on there being high confidence that the MAVE assay relates to the mechanism of disease. For each specific gene-disease relationship in question, ensuring that the MAVE assay is a valid predictor of pathogenicity will be essential. This is likely to be especially challenging for genes with multiple different disease associations and for proteins with multiple functional domains. We believe that the mapping of these data to human reference sequence assemblies provides a crucial foundation for the development of AVE guidelines for the use of MAVE data, by providing common sequence assemblies for the evaluation of MAVE score sets.

## Conclusions

The impact of genome and exome sequencing on human research and clinical practice is hindered by challenges in variant interpretation. Multiplexed assays of variant effect (MAVEs) provide a high-throughput functional assessment tool for variants in genes of relevance to human health and disease, and hundreds of MAVEs have been developed and submitted to the centralized MaveDB data repository. We developed a method for precise representation and mapping of MAVE data to human reference sequences and application of this method to score sets in MaveDB. We demonstrate how this process enables the mapping of 98.61% (2,464,212 / 2,499,044) of MAVE variants in MaveDB for ready integration into multiple variant annotation and evaluation platforms. We discuss current challenges in the use of these data in human genomics applications, and the need for expert communities like the Atlas of Variant Effects Alliance (AVE) to address these remaining gaps. We believe the mapped data from this study will help advance those efforts, and the data integrations at the UCSC Genome Browser, Genomics 2 Proteins Portal, Ensembl VEP, ClinGen Linked Data Hub, and others will provide useful tools for advancing MAVE-informed genomic variant interpretation efforts.

## Methods

### Extraction of Metadata from the MaveDB API Score Sets Endpoint

For score sets whose listed target organism was *Homo sapiens*, seven variables were extracted from the MaveDB API score sets endpoint. These variables were: target sequence (string of nucleotides or amino acids), target sequence type (DNA or protein), target (e.g. *CXCR4*), assembly ID, UniProt ID, target type (e.g. protein coding), and URN (e.g. urn:mavedb:00000048-a-1). These data elements were extracted from the MaveDB API as they were the minimum information needed to determine the genomic coordinates targeted by an assay in a MaveDB score set.

### Alignment of Target Sequences to Human Genome

Having extracted the necessary metadata from the MaveDB API, the initial step was to align target sequences to the human genome, allowing for the genomic coordinates of the examined sequences to be determined (**Figure 4**). To achieve this aim, all target sequences were run through BLAT against the GRCh38 human genome assembly. Depending on whether a target sequence was composed of nucleotides or amino acids, the BLAT query (q) argument was set to “dna” or “prot” to maximize the probability of returning high quality hits. In addition, the minimum score argument (minScore) was reduced to 20 from a default value of 30 to ensure that BLAT would return hits from short, specific target sequences.

After running BLAT, a series of steps were followed to ensure that suitable genomic coordinates were located. When running BLAT locally and outputting the results in a Pattern Space Layout (PSL) file, different “hits’’ are reported at the chromosomal level (e.g. “chr3”). Within a hit are specific “HSP” (high-scoring pair) objects that describe regions of concordance between the queried sequence and the human genome. With the goal of ultimately selecting the correct HSP object, the correct chromosomal hit needs to be chosen first. To maximize this probability, when a UniProt accession was available for a score set, the UniProt accession was supplied as a compact uniform resource identifier (CURIE, e.g. uniprot:P12931) to the normalization method in the Variant Interpretation for Cancer Consortium (VICC) Gene Normalization Service(35), returning an HGNC(36) consensus gene symbol and a chromosome number indicating where the gene occurs. Based on the chromosome number returned from the normalize method, the BLAT PSL file was filtered to only include the hit that contained the correct chromosome. If the filtered hit contained more than one HSP object, additional processing was required to select the correct genomic coordinates. First, the gene symbol corresponding to the provided UniProt accession was supplied to the search method within the Gene Normalization Service, returning location data, in genomic coordinates, from Ensembl(37) and NCBI(38). Specifically, these two sources had “start” and “end” attributes describing the location where the supplied gene of interest occurs on a chromosome. Using the position indicated in the start attribute, the HSP object with the minimum distance to the start position was selected.

Having performed a series of filtering and validation steps to ensure that the genomic coordinates with the highest potential accuracy were selected, alignment data was added to a dictionary that was keyed by score set URN. For each score set, the supplied data included: chromosome number, strand orientation, target name, target type, UniProt ID, percent coverage, percent identity, and a dataframe reporting the genomic coordinates supplied by the selected HSP object.

While the alignment procedure performs well for protein coding score sets with UniProt accessions, its efficacy is potentially limited for regulatory/other noncoding score sets. Specifically, these score sets lack UniProt accessions and often have more descriptive target names without gene names (e.g. hYAP65 WW domain); the ability to extract consensus gene symbols for these score sets is limited, thereby resulting in a potential inability to perform the hit/HSP filtering procedure. In these instances, the top scoring hit reported by BLAT was selected, but the additional validation steps described previously were not performed.

### RefSeq Transcript Selection and Offset Computation

Having extracted genomic coordinates and other relevant information such as the chromosome number, there was sufficient data for selecting an appropriate human reference sequence for each score set. For the regulatory/other noncoding score sets that only reported genomic variation, the RefSeq chromosomal genomic reference sequence for GRCh38 was selected since the supplied genomic coordinates described locations on the sequence. However, for protein coding score sets that reported protein variants, a RefSeq protein reference sequence was needed, requiring use of the Biocommons UTA and SeqRepo databases and the associated Cool-Seq-Tool(30) translation service.

First, using SeqRepo, a database that stores human reference sequences and links between different identifiers, chromosome numbers were converted to their GRCh38 RefSeq accession. Additionally, consensus gene symbols were derived by leveraging the Gene Normalization service that was utilized in the alignment procedure. Lastly, a query was run against UTA that extracted all the transcripts found within the genomic start and end coordinates, supplied by the HSP fragment, that were associated with the derived chromosome accession and gene symbol, and stored the identifiers in a list. This query was run for all rows in the HSP dataframe; for example, if an HSP object from a score set reported six fragments, six lists were generated. Once all lists were created and any non-coding transcript accessions were removed, the intersection of the lists was taken and supplied as input to the get_mane_from_transcripts() method in Cool-Seq-Tool. If the method returned a nonempty list, the following prioritization ranking was followed for transcript selection: MANE Select, MANE Plus Clinical. If the list was empty, the lengths of all transcripts in the intersected list were taken, the longest transcript was chosen (with the first-published remaining transcript breaking ties), and the protein RefSeq accession associated with the selected transcript was found.

To determine the exact location of the provided target sequence within the RefSeq protein sequence, the target sequence, if DNA, was converted to protein using the standard codon table and the first 10 amino acids were extracted. Then, by accessing the RefSeq sequence using SeqRepo and using the find() method, the substring’s location was found; find() was run again a second time on the entire converted protein sequence, producing a boolean that indicated if the *entire* target sequence was an exact substring of the RefSeq sequence. When the intersection procedure returned a non-RefSeq sequence, web scraping was performed to extract the canonical UniProt sequence and find() was again run twice. Lastly, the protein reference sequence, offset, score set accession, transcript and MANE status, and boolean were stored as entries in a dictionary and saved to a pickle file.

After an initial offset was computed for each protein coding score set, an additional pass through the score sets was completed to check for possible discordance between the provided target sequences and corresponding variant matrices. Assume a target sequence is a string of amino acid residues and reports Leucine at position 100. If the score matrix reports a substitution such as “p.Arg100His,” this would be an example of discordance as the expected reference amino acid would be Leucine instead of Arginine. To identify instances of such discordance, for each score set, the provided target sequence, if DNA, was first converted to a protein sequence using the standard codon table. Then, the variant matrix was parsed to assemble a dictionary reporting the expected reference amino acid at each position (ex. 1: M, 2: A, etc.). By comparing the expected amino acids from the dictionary to those in the corresponding positions in the target sequence, the correct start location in the target sequence was determined and the offset was modified to reflect this position.

The general offset computed by the described procedure was applicable for protein variants in protein coding score sets. For genomic variants in protein coding score sets, the correct mapped position was found by determining the alignment block that the variant was found in and computing the distance between the variant position and the start (for positive strand) or end (for negative strand) genomic coordinate of the selected block, depending on the orientation of the target sequence. The same logic was followed for regulatory/other noncoding score sets with genomic variants that had only one alignment block.

### Mapping Variants using VRS

With the relevant human reference sequence data determined, variant locations reported in MaveDB score set matrices were updated and the variants themselves were expressed as VRS objects. First, for a given row in a score set, the variant was first converted to a VRS object as is, allowing for the VRS representation of the assayed variant to be generated. Specifically, the reported positions were left unchanged, a new sequence digest was computed using the sha512t24u digest algorithm, the allele was normalized using the SPDI Variant Overprecision Correction Algorithm (VOCA)(39), and the appropriate allele digest was determined using the VRS ga4gh_identify() method. In instances where multiple variants were indicated with the semicolon character, this process was run separately for each variant and a VRS haplotype was generated. When a variant type not supported by VRS appeared, although a computable identifier could not be assigned, a VRS text variation was generated. All processed pre-mapped and post-mapped variants were represented using VRS version 1.3.

The process described above was then repeated using the information derived from the reference sequence selection procedure. Using the translate_identifier() method in SeqRepo, the sequence digest was updated to describe the digest for the human reference sequence while the offset was added to start and end position values, respectively. A new allele digest was computed, allowing for the “mapped” variant to have a distinct identifier from the “assayed” variant. When multiple variants were reported, new VRS alleles were created for each variant and combined in a VRS haplotype. The assayed and mapped variant representations for each row were then converted to dictionaries, allowing for storage in a two-column dataframe.

It is possible that an individual uploading a score set to MaveDB could choose to represent variants using a human reference sequence. For example, score sets in experiment set urn:mavedb:00000097 report variants in the “hgvs_prot” column with a RefSeq protein accession (e.g. NP_009225.1:p.Pro1659Leu). When this occurred, the variant was directly converted to a VRS allele using the translate_from() method and the assayed and mapped representations of the variant were equivalent.

As the regulatory/other noncoding score sets only reported variants in the “hgvs_nt” column, one set of VRS representations was produced for each of these score sets. However, as protein coding score sets were capable of having data in both the “hgvs_nt” column *and* “hgvs_prot” column, variants in both columns, if present, were mapped, resulting in two potential sets of mappings per score set. Once mappings were complete for all score sets, a dictionary, keyed by score set URN and containing the mappings for protein coding and regulatory/other noncoding score sets, was generated and saved as a pickle file.

### Annotating and Validating VRS Alleles/Haplotypes Reference Sequences

The pickle files containing the assayed and mapped variants were further annotated with the “vrs_ref_allele_seq” attribute that allows for the reference allele for a VRS object to be reported along with the modified allele. To ensure that all VRS objects in all processed score sets contained this attribute, a VRS object was instantiated for each pre-mapped and post-mapped variant, where the existing allele digest was supplied to the “id” attribute and the VRS allele was supplied to the “variation” attribute. SeqRepo was utilized to determine the reference allele sequence for the pre-mapped and post-mapped objects and the sequence was accordingly provided to the “vrs_ref_allele_seq” attribute. Additionally, for post-mapped variants, an HGVS string (e.g. “NC_000006.12:g.37808023C>A”) describing the VRS allele was generated and supplied to the “expressions’’ attribute in post-mapped alleles.

The completion of this step validated the accuracy of the mapping procedure as it provided a check on the consistency between the reference alleles generated for the pre-mapped and post-mapped representations. For protein variants and genomic variants derived from positively oriented target sequences, we observed concordance between substrings provided by the “vrs_ref_allele_seq” attribute for the pre-mapped and post-mapped objects. However, as SeqRepo contains reference sequences derived from positive strands, the reference allele sequence was the reverse complement for genomic variants that came from negatively oriented target sequences (e.g. pre-mapped: “GAT”, post-mapped: “ATC”).

### Mapping File Format

The newly created pre-mapped and post-mapped objects were added to a “mapped_scores’’ attribute for each score set. Additionally, a “computed_reference_sequence” attribute, storing the target sequence, sequence type, and sequence digest, and a “mapped_reference_sequence” attribute, storing the RefSeq accession, sequence type, and corresponding sequence digest, were added to each score set. Following the creation of the respective attributes, all processed score sets were saved as JSON files, gzipped, and uploaded to a publicly-accessible s3 bucket (mavedb-mapping).

### Integrating MaveDB Data into the UCSC Genome Browser

Due to the large amount of protein-level variation data reported by MAVEs, additional tools needed to be employed to translate protein changes to codons. This was accomplished with the VICC protein to genome mapping method from Cool-Seq-Tool (30) at https://normalize.cancervariants.org/variation/alignment_mapper/p_to_g to map the protein-level variants to the genomic coordinates associated with their corresponding codons. Given these genomic coordinates, the MAVE score sets were translated visually to heat maps using the UCSC Genome Browser’s bigHeat utility (https://hpc.nih.gov/apps/Genome_Browser.html): this utility merges the genomic coordinates, contained in an input BED file, with a “location matrix” (a TSV file that reports scores per bed item), and generates one bigBed file per column, colored by the values in the matrix. This yielded a set of Genome Browser tracks, each of which represent one MaveDB score set.

### Integrating MaveDB Data into the ClinGen LDH

To integrate the MaveDB data into the ClinGen LDH, a MaveDBMapping document was created for each score set entry in the mapping files and added to the LDH as linked data for an LDH variant represented by the ClinGen Allele Registry canonical allele identifier. Because the ClinGen Allele Registry requires the use of standard human reference sequences (genome builds NCBI36, GRCh37, GRCh38 and transcripts from NCBI or Ensembl), each HGVS expression within the post-mapped objects from these score set entries was leveraged to either find the existing canonical allele identifier referenced in the score set entry or to register the variant with the ClinGen Allele Registry to obtain a new canonical allele identifier. MaveDbMapping documents were created by excerpting the MaveDB mapped scores object, score, MaveDB score set id (URN + entry number; eg. urn:mavedb:00000001-a-1#1), captured provenance information (creation, modification and publish dates), and a link back to the referenced MaveDB score set page.

## Abbreviations

API: Application Programming Interface
AVE: Atlas of Variant Effects Alliance
BLAT: BLAST-like Alignment Tool
CAid: Canonical Allele Identifier
Cool-Seq-Tool: Common Operations on Lots of Sequences Tool
CURIE: Compact Uniform Resource Identifier
DCD: Data Coordination and Dissemination Workstream
DECIPHER: Database of Genomic Variation and Phenotype in Humans using Ensembl Resources
G2P: Genomics 2 Proteins Portal
GA4GH: Global Alliance for Genomics and Health
HGNC: Human Gene Nomenclature Committee
HGVS: Human Genome Variation Society
HSP: High-Scoring Pair
JSON: JavaScript Object Notation
LDH: ClinGen Linked Data Hub
MANE: Matched Annotation from NCBI and EMBL-EBI
MAVE: Multiplexed Assays of Variants Effect
MPRA: Massively Parallel Reporter Assay
NCBI: National Center for Biotechnology Information
PAid: Protein Allele Identifier
PSL: Pattern Space Layout
PyPi: Python Package Interface
RefSeq: NCBI Reference Sequence Database
SO: Sequence Ontology
URN: Uniform Resource Name
UTA: Universal Transcript Archive
VEP: Ensembl Variant Effect Predictor
VICC: Variant Interpretation for Cancer Consortium
VOCA: Variant Precision Overcorrection Algorithm
VRS: Global Alliance for Genomics and Health Variation Representation Specification
VUS: Variant of Uncertain Significance

## Declarations

### Ethics approval and consent to participate

Not applicable

### Consent for publication

Not applicable

### Availability of data and materials

The code and datasets generated during this study are publicly available under the MIT license in the dcd_mapping repository at: https://github.com/ave-dcd/dcd_mapping. The source code used in this study is stably archived using Zenodo at https://zenodo.org/records/11406658.

### Authors’ Contributions

AFR and AHW conceptualized the study. JAA, SI, MSC, AFR, and AHW developed the methodology. JAA, EYD, JSS, KK, TF, BJC, SBG, KR, NSA, AM, SEH, MSC, AFR, and AHW developed software. JAA and SI performed validation. JAA, JFS, and SI performed formal analysis. JAA, JFS, and SI conducted investigations. MSC provided resources. AP curated data. JAA, JSS, JFS, SI, AFR, and AHW wrote the original draft of the manuscript. JAA, JSS, KK, TF, KR, JFS, JF, HVF, SEH, SI, AFR, and AHW revised and edited the manuscript. JAA, JFS, and SI performed data visualization. SEH, SI, AFR, and AHW supervised the project. AFR and AHW performed project administration. All authors read and approved the final manuscript.

## Acknowledgements

We would like to thank Dr. William Ray and the Graphics Services team at Nationwide Children’s Hospital for their assistance in developing the figures for this manuscript.

