## Supplementary figures and images for "Mapping MAVE data for use in human genomics applications"

### Supplementary Figure 1

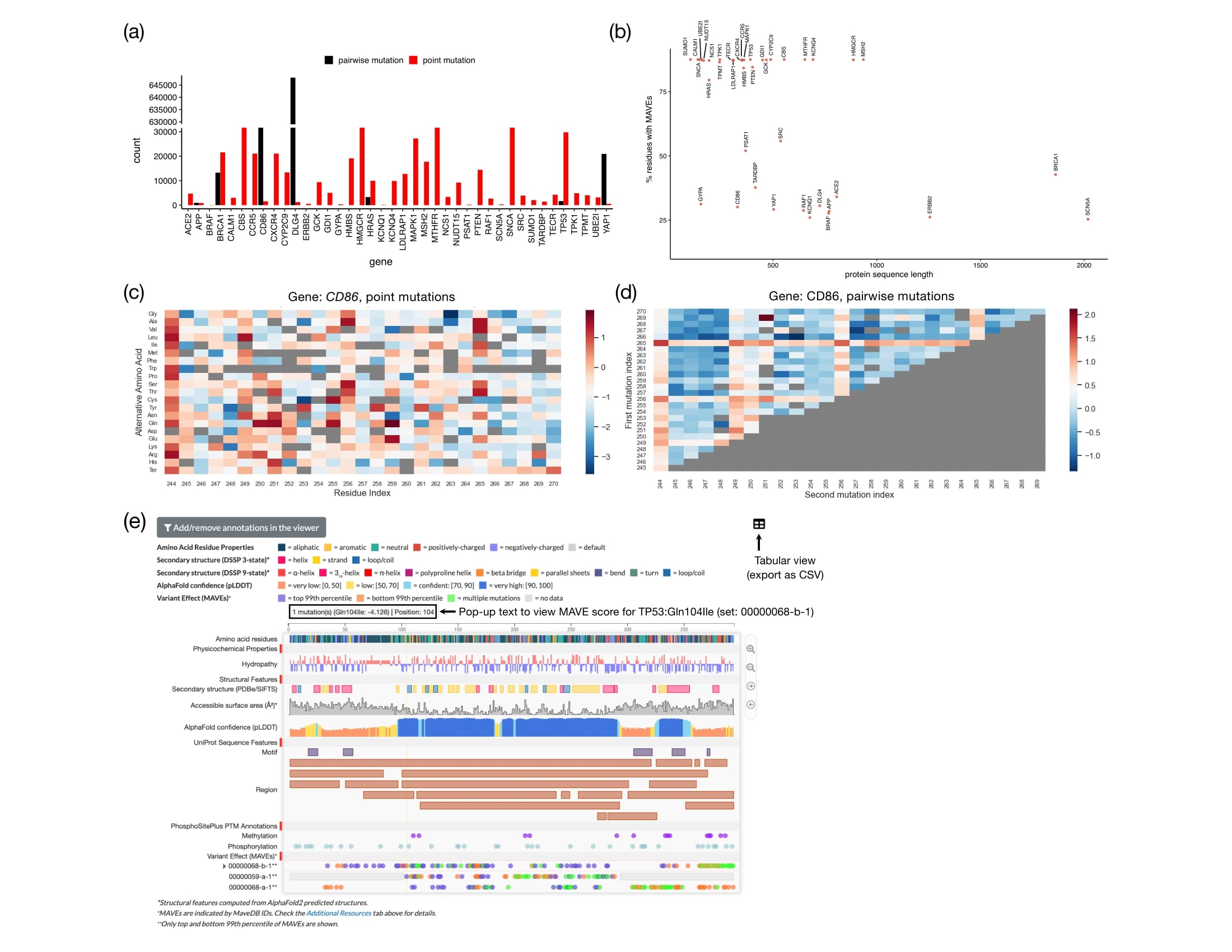

### Supplementary Figure 2

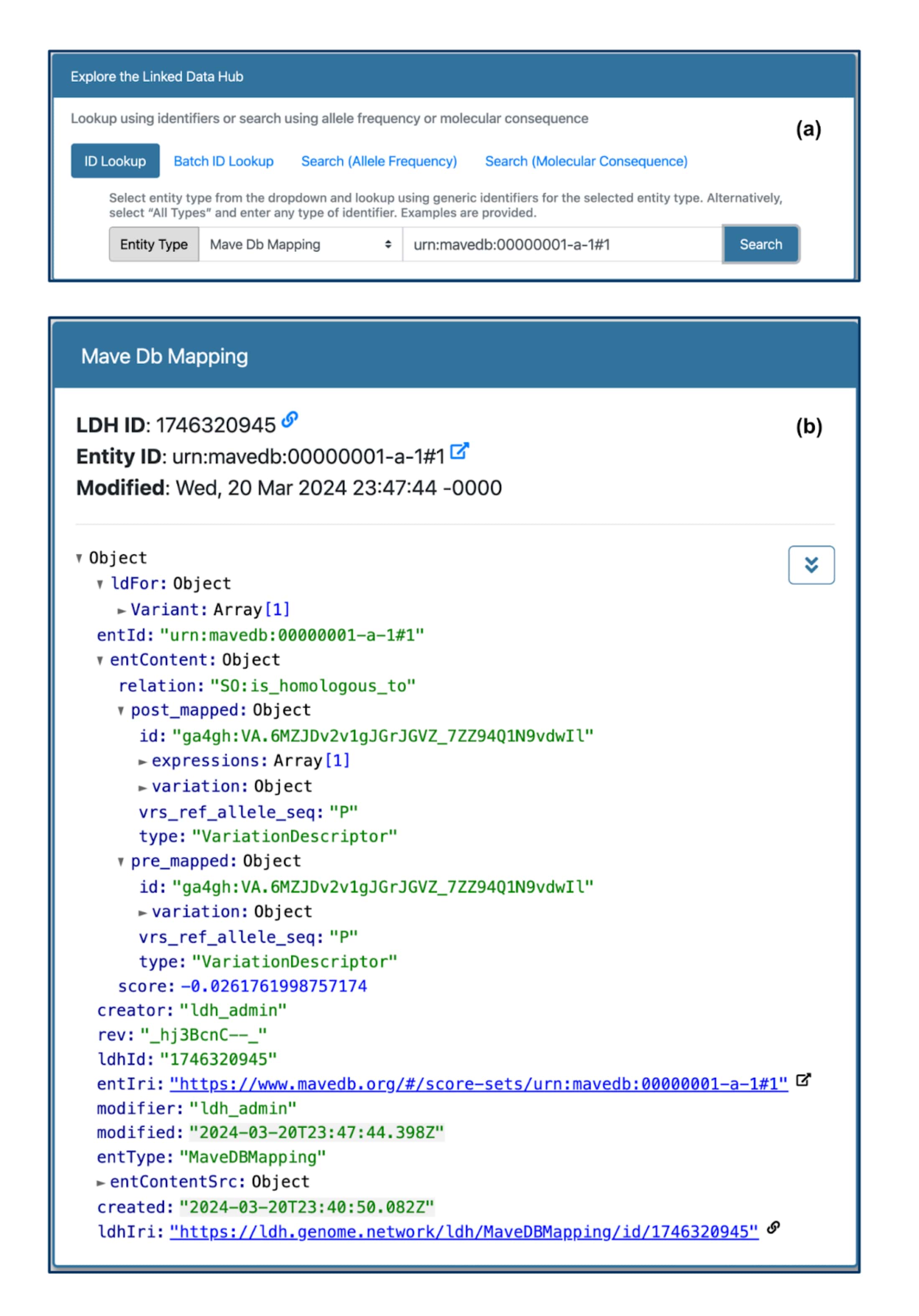
